# Inducible deletion of *Ezh2* in CD4+ T cells inhibits kidney T cell infiltration and prevents interstitial nephritis in MRL/*lpr* lupus-prone mice

**DOI:** 10.1101/2024.03.04.583401

**Authors:** Xiaoqing Zheng, Mikhail G Dozmorov, Luis Espinoza, Mckenna M Bowes, Sheldon Bastacky, Amr H Sawalha

## Abstract

Systemic lupus erythematosus is a remitting relapsing autoimmune disease characterized by autoantibody production and multi-organ involvement. T cell epigenetic dysregulation plays an important role in the pathogenesis of lupus. We have previously demonstrated upregulation of the key epigenetic regulator EZH2 in CD4+ T cells isolated from lupus patients. To further investigate the role of EZH2 in the pathogenesis of lupus, we generated a tamoxifen-inducible CD4+ T cell *Ezh2* conditional knockout mouse on the MRL/*lpr* lupus-prone background. We demonstrate that *Ezh2* deletion abrogates lupus-like disease and prevents T cell differentiation. Single-cell analysis suggests impaired T cell function and activation of programed cell death pathways in EZH2-deficient mice. *Ezh2* deletion in CD4+ T cells restricts TCR clonal repertoire and prevents kidney-infiltrating effector CD4+ T cell expansion and tubulointerstitial nephritis, which has been linked to end-stage renal disease in patients with lupus nephritis.

## Introduction

Systemic lupus erythematosus (SLE or lupus) is a heterogenous remitting-relapsing autoimmune disease that can affect multiple organ systems. Lupus is characterized by autoantibody production, type-I interferon gene expression signature, and DNA methylation defect (1). Indeed, abnormal T cell DNA methylation is thought to play a central role in the pathogenesis of lupus, in part due to a role for DNA demethylation in inducing T cell autoreactivity (2). Further, lupus T cells are characterized by robust demethylation of interferon-regulated genes, which predates T cell activation in lupus patients (3).

We have previously demonstrated an epigenetic shift in lupus CD4+ T cells that occurs upon increased disease activity. As the disease becomes more active, naïve CD4+ T cells are shifted into epigenetic patterns more favorable for proinflammatory T helper cell subsets, and away from differentiation into regulatory T cells (4). These epigenetic changes occur early in the naïve CD4+ T cell stage and precede corresponding transcriptional changes or T cell differentiation. Bioinformatics analysis followed by experimental work suggested that the key epigenetic regulator EZH2 might be underlying the observed disease activity-associated CD4+ T cell epigenetic changes (4, 5). Indeed, we were able to demonstrate increased expression of EZH2 in lupus CD4+ T cells, and that EZH2 overexpression in normal CD4+ T cells leads to proinflammatory epigenetic changes, such as demethylation and overexpression of the adhesion molecule JAM-A (5). Our work further demonstrated that reduced expression of two microRNAs that regulate EZH2 (miR-101 and miR-26a) underlies EZH2 overexpression in lupus T cells (5). We further demonstrated that increased glycolysis in lupus T cells explains reduced expression of mi-R101 and miR-26a, and that inhibiting glycolysis, either directly or indirectly via blocking mTOR signaling, restores miR-101 and miR-26a levels and inhibits the expression of EZH2 (6).

To demonstrate a pathogenic role for EZH2 in lupus, we treated MRL/*lpr* lupus-prone mice with an EZH2 inhibitor and showed reduced autoantibody production and amelioration of lupus-like disease (7). To further elucidate the role of EZH2 in lupus CD4+ T cells specifically, we herein generated an MRL/*lpr* mouse with an inducible genetic deletion of *Ezh2* in CD4+ T cells. Using a preventative and a therapeutic model, we show that EZH2 deficiency in CD4+ T cells is sufficient to prevent autoantibody production and improve lupus-like manifestations. We further show that T cell-EZH2 deficiency prevents activation of naïve CD4+ T cells and reduces kidney infiltrating activated T cells as well as interstitial nephritis in MRL/*lpr* mice.

## Results

### Tamoxifen-inducible CD4+ T cell-specific deletion of *Ezh2* in MRL/*lpr* mice

We generated a tamoxifen-inducible CD4+ T cell-specific EZH2-deficient mouse (iCD4-Cre *Ezh2^fl/fl^*) on the MRL/*lpr* lupus-prone background as described above. *Ezh2^fl/fl^* mice without iCD4-Cre were used as controls. Tamoxifen induced *Ezh2* deletion in iCD4-Cre *Ezh2^fl/fl^* compared to *Ezh2^fl/fl^* control mice (**Figure 1A**). No deletion of *Ezh2* was detected in other cell types, including CD8+ T cells (data not shown).

**Figure 1.**
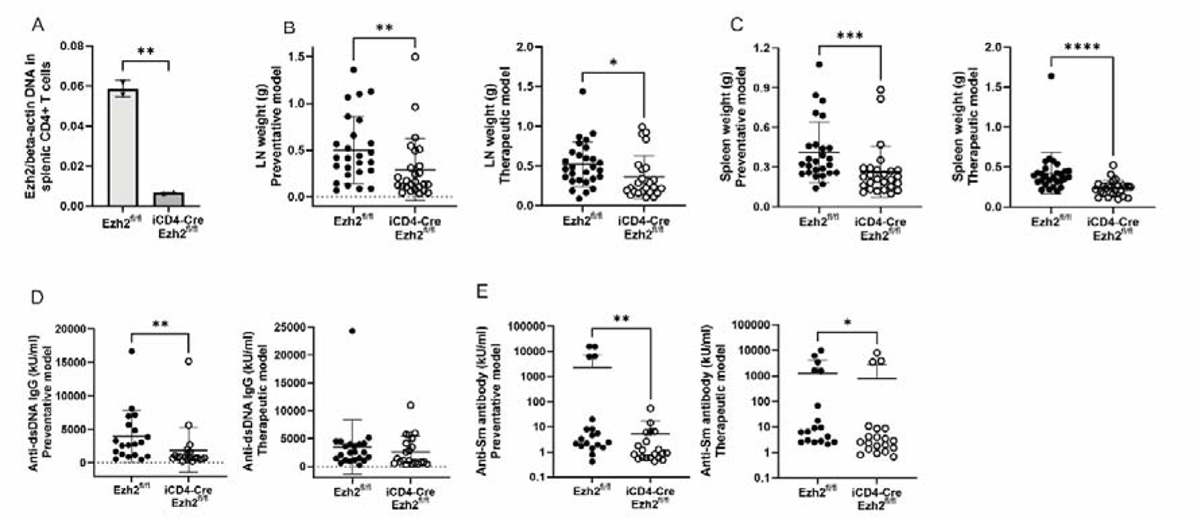
(**A**) *Ezh2* DNA level in CD4+ T cells isolated from *Ezh2^fl/fl^* and CD4-Cre *Ezh2^fl/fl^* mice after 300 mg/kg tamoxifen treatment for 3 days (Monday, Wednesday, and Friday) by oral gavage. Data are shown as mean ± SD. ** *p*<0.01 (n=2). (**B**) Weight of cervical lymph nodes in *Ezh2^fl/fl^* and iCD4-Cre *Ezh2^fl/fl^* mice from the preventative and therapeutic mouse models. LN: lymph nodes; results are presented as mean ± SD, \**p*<0.05, \*\**p*<0.01, two-tailed Mann-Whitney test. (**C**) Spleen weight in *Ezh2^fl/fl^* and iCD4-Cre *Ezh2^fl/fl^* mice from the preventative and therapeutic mouse models. Data are presented as mean ± SD, \*\*\**p*<0.001, \*\*\*\**p*<0.0001, two-tailed Mann-Whitney test. (**D, E**) Anti-dsDNA IgG and anti-Sm antibody levels in the preventative and therapeutic mouse models. Anti-dsDNA: anti-double stranded DNA. Results are presented as mean ± SD, \**p*<0.05, \*\**p*<0.01, two-tailed Mann-Whitney test.

### *Ezh2* deletion in CD4+ T cells alleviated lupus-like disease

*Ezh2* deletion in CD4+ T cells significantly inhibited peripheral lymphoproliferation, as evidenced by reduced weight of cervical lymph nodes and the spleen in iCD4-Cre *Ezh2^fl/fl^* compared to *Ezh2^fl/fl^* control mice in preventative and therapeutic models (**Figure 1B and 1C**). These differences were observed in both female and male mice (**Supplementary** Figure 1).

The production of anti-dsDNA IgG autoantibodies was significantly reduced in EZH2 deficient mice in the preventative but not the therapeutic model (**Figure 1D**). Anti-Sm autoantibodies were significantly lower in iCD4-Cre *Ezh2^fl/fl^* compared to littermate *Ezh2^fl/fl^* controls in both models (**Figure 1E**).

We examined the effect of CD4+ T cell-*Ezh2* deletion on lupus nephritis in MRL/*lpr* mice. Urine albumin to creatinine ratio (UACR) was significantly reduced in both the preventative and therapeutic models (**Figure 2A**). Kidney tissues were fixed, embedded in paraffin, sections were stained with hematoxylin and eosin (H&E), and a renal pathologist reviewed the slides in a blinded manner. Scores were given based on inflammatory changes in the glomeruli and interstitial areas as previously described (8). EZH2 deficiency in CD4+ T cells did not affect glomerular inflammation (**Figure 2B**). However, interstitial nephritis was significantly reduced in CD4+ T cell EZH2-deficient mice compared to controls (**Figure 2C**). Representative kidney H&E staining images from iCD4-Cre *Ezh2^fl/fl^* and *Ezh2^fl/fl^* control mice are shown in **Figure 2D**.

**Figure 2.**
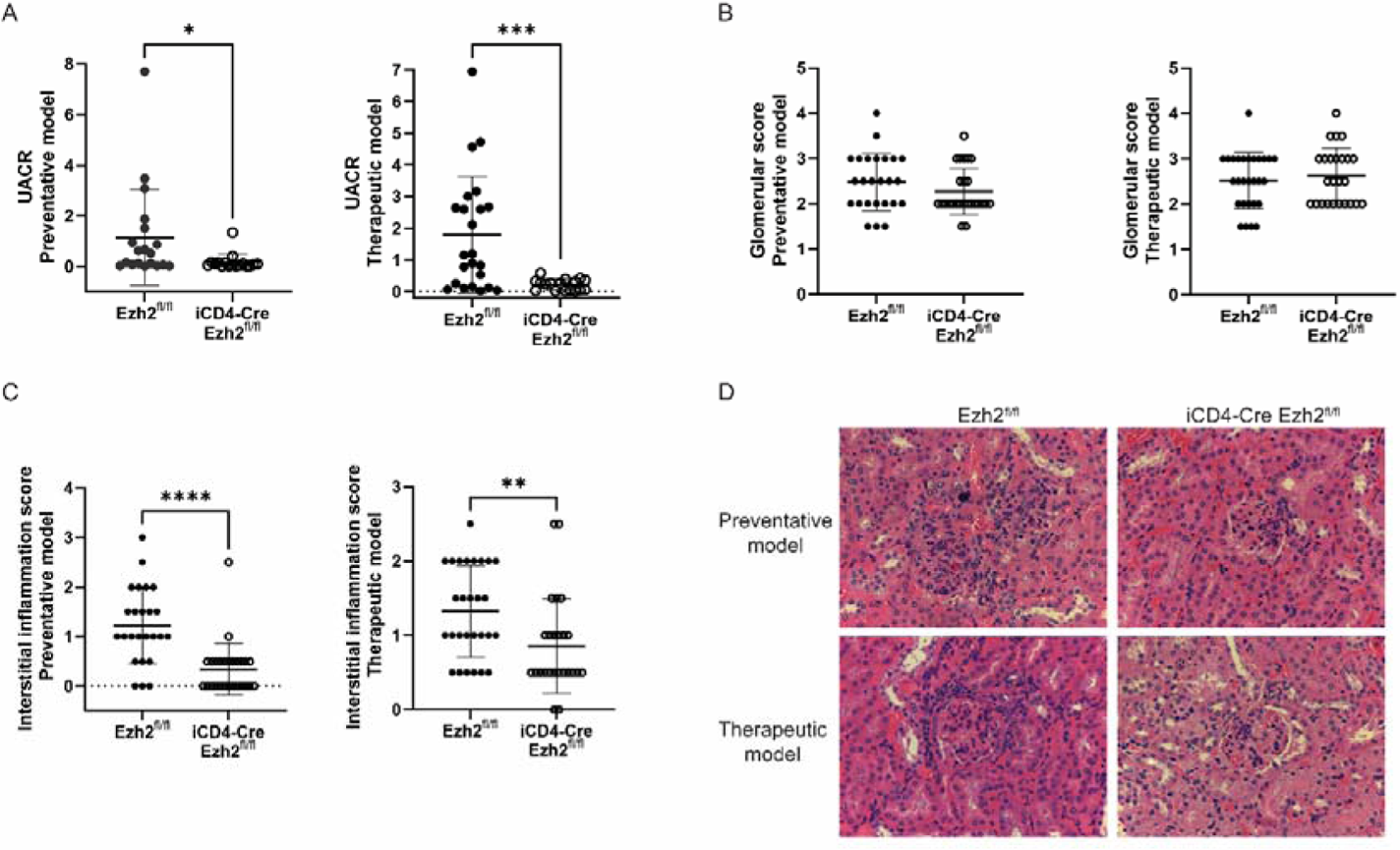
(**A**) Urine albumin to creatinine ratio (UACR) in the preventative and therapeutic mouse models. Data are presented as mean ± SD. \**p*<0.05, \*\*\**p*<0.001, two-tailed Mann-Whitney test. Glomerulonephritis scores (**B**) and interstitial nephritis scores (**C**) in the preventative and therapeutic mouse models. Data are presented as median with interquartile range, ** *p*<0.01, \*\*\*\**p*<0.0001, two-tailed Mann-Whitney test. (**D**) Representative hematoxylin and eosin (H&E) images of kidney tissues. Original magnification: 400x.

### EZH2 deficiency impedes CD4+ T cells activation and differentiation

We observed a significant reduction in CD4+ T cells in the spleen in iCD4-Cre *Ezh2^fl/fl^* compared to *Ezh2^fl/fl^* control mice in the preventative model (**Figure 3A and 3B**). While there was an increase in the percentage of CD8+ T cells in iCD4-Cre *Ezh2^fl/fl^* mice, there was no significant change in the number of CD8+ T cells in the spleen (**Figure 3C**). No significant difference was observed in double negative (CD3ε+ TCRβ+CD4-CD8-) T cells (**Figure 3D**). Gating strategies are shown in **Supplementary** Figure 2.

**Figure 3.**
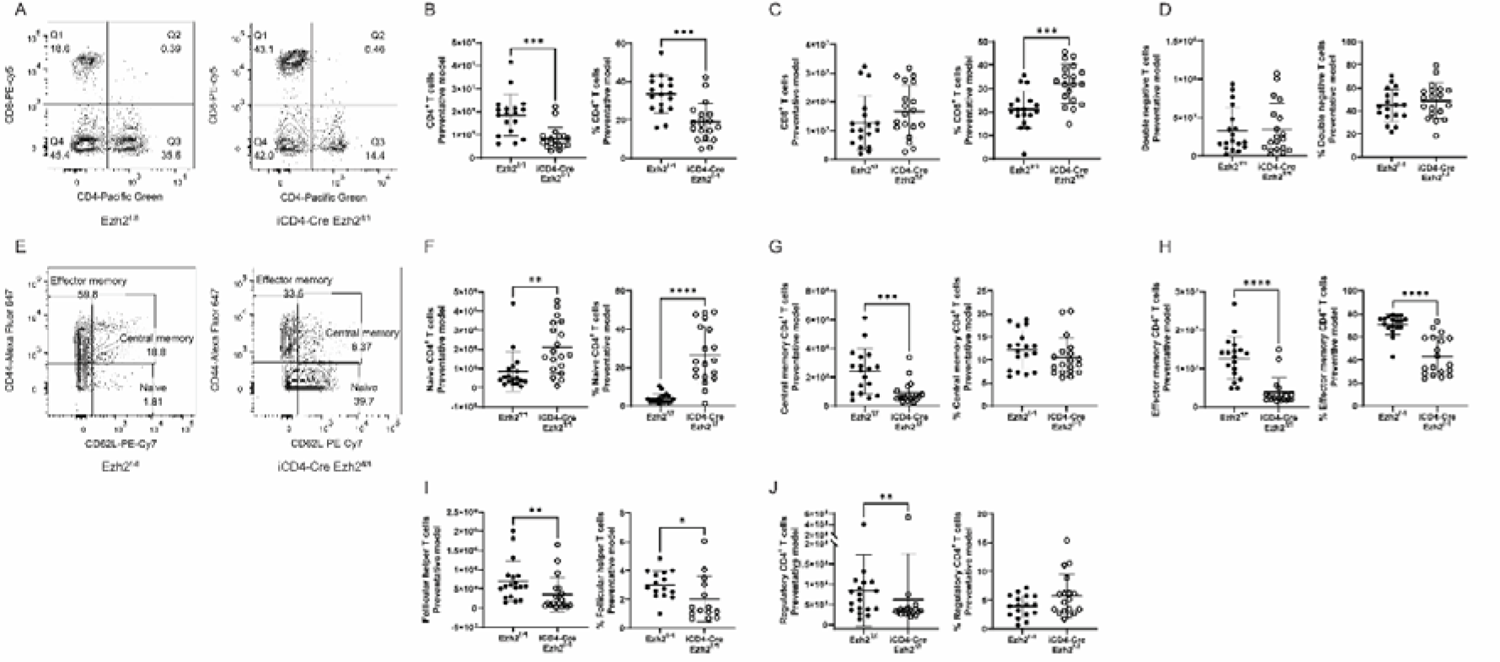
Representative flow cytometry plot (**A**) and statistical analysis of CD4+ T cell (**B**), CD8+ T cells (**C**) and double negative (CD4-CD8-) T cells (**D**) in the spleen of *Ezh2^fl/fl^* and iCD4-Cre *Ezh2^fl/fl^* mice. Representative flow cytometry plot (**E**) and statistical analysis of naïve CD4+ T cells (**F**), central memory CD4+ T cells (**G**) and effector memory CD4+ T cells (**H**) in the spleen of *Ezh2^fl/fl^* and iCD4-Cre *Ezh2^fl/fl^* mice. Statistical analysis of follicular helper T cells (**I**) and regulatory CD4+ T cells (**J**) in the spleen of *Ezh2^fl/fl^* and iCD4-Cre *Ezh2^fl/fl^* mice. Data are presented as mean ± SD. \**p*<0.05, \*\**p*<0.01, \*\*\**p*<0.001, \*\*\*\**p*<0.0001, two-tailed Mann-Whitney test.

Among CD4+ T cells in the spleen, the frequency and numbers of naïve CD4+ T cells was significantly higher in iCD4-Cre *Ezh2^fl/fl^* compared to *Ezh2^fl/fl^* control mice, and CD4+ T cell differentiation into effector memory CD4+ T cells was significantly inhibited with EZH2 deficiency. Indeed, the majority of CD4+ T cells were naïve T cells in iCD4-Cre *Ezh2^fl/fl^* mice, while there were activated and differentiated into effector memory and central memory CD4+ T cells in *Ezh2^fl/fl^* control mice (**Figure 3E-H**). We also observed a reduction in the number of follicular helper T cells and regulatory CD4+ T cells in EZH2-deficient mice (**Figure 3I and 3J**). Similar data were obtained from the therapeutic mouse model (**Supplementary** Figure 3).

### EZH2 deficiency inhibits renal infiltration of CD4+ T cells

We evaluated kidney-infiltrating T cells with and without EZH2 deficiency in the preventative model. We found a significant reduction in kidney-infiltrating T cells in iCD4-Cre *Ezh2^fl/fl^* compared to control mice, explained by a reduction in CD4+ T cells (**Figure 4A-F**). Similar to the spleen, kidney-infiltrating CD4+ T cells from iCD4-Cre *Ezh2^fl/fl^* mice had significantly fewer central memory and effector memory CD4+ T cells compared to littermate *Ezh2^fl/fl^* control mice (**Figure 4G-J**).

**Figure 4.**
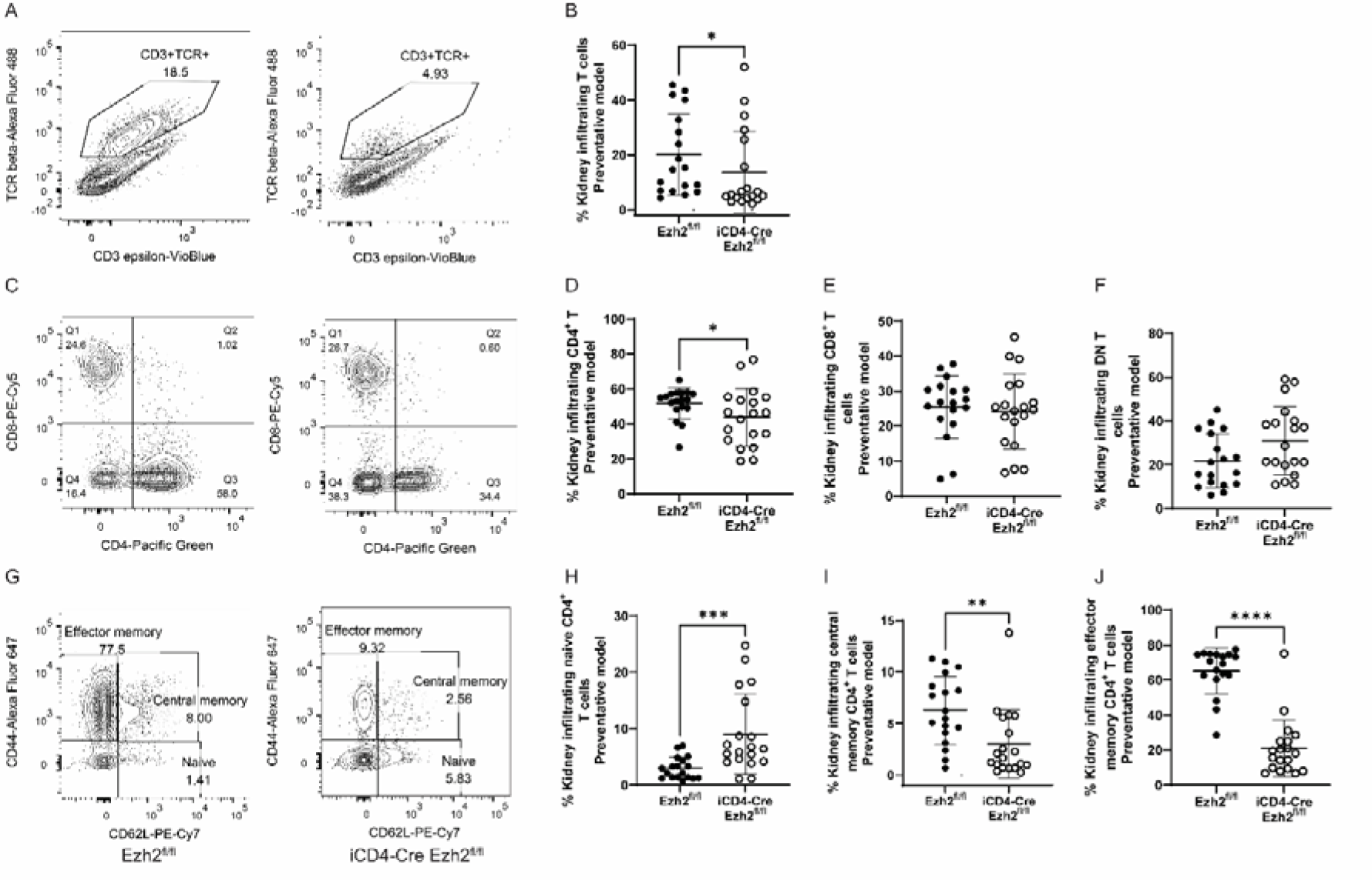
Representative flow cytometry plot (**A**) and statistical analysis of (**B**) kidney infiltrating CD3ε+ TCRβ+ T cells in *Ezh2^fl/fl^* and iCD4-Cre *Ezh2^fl/fl^* mice. Representative flow cytometry plot (**C**) and statistical analysis of kidney infiltrating CD4+ T cells (**D**), CD8+ T cells (**E**) and double negative (DN, CD4-CD8-) T cells (**F**) in *Ezh2^fl/fl^* and iCD4-Cre *Ezh2^fl/fl^* mice. Representative flow cytometry plot (**G**) and statistical analysis of kidney infiltrating naïve CD4+ T cells (**H**), central memory CD4+ T cells (**I**) and effector memory CD4+ T cells (**J**). Data are presented as mean ± SD, \**p*<0.05, ** *p*<0.01, \*\*\**p*<0.001, \*\*\*\**p*<0.0001, two-tailed Mann-Whitney test.

### Single-cell RNA-seq reveals T cell subset compositional changes with *Ezh2* deletion

To further explore the effects of EZH2 deficiency in CD4+ T cells, single-cell RNA sequencing was performed in FACS-sorted CD4+ T cells isolated from the spleen. In total, 15 cell populations were clustered based on cell transcriptional profiles. Cell identity of each cluster was determined by evaluating differentially expressed genes within the clusters. We annotated each cluster as demonstrated in **Figure 5A**, 2 of which were non-T cell clusters (monocytes and macrophages).

**Figure 5.**
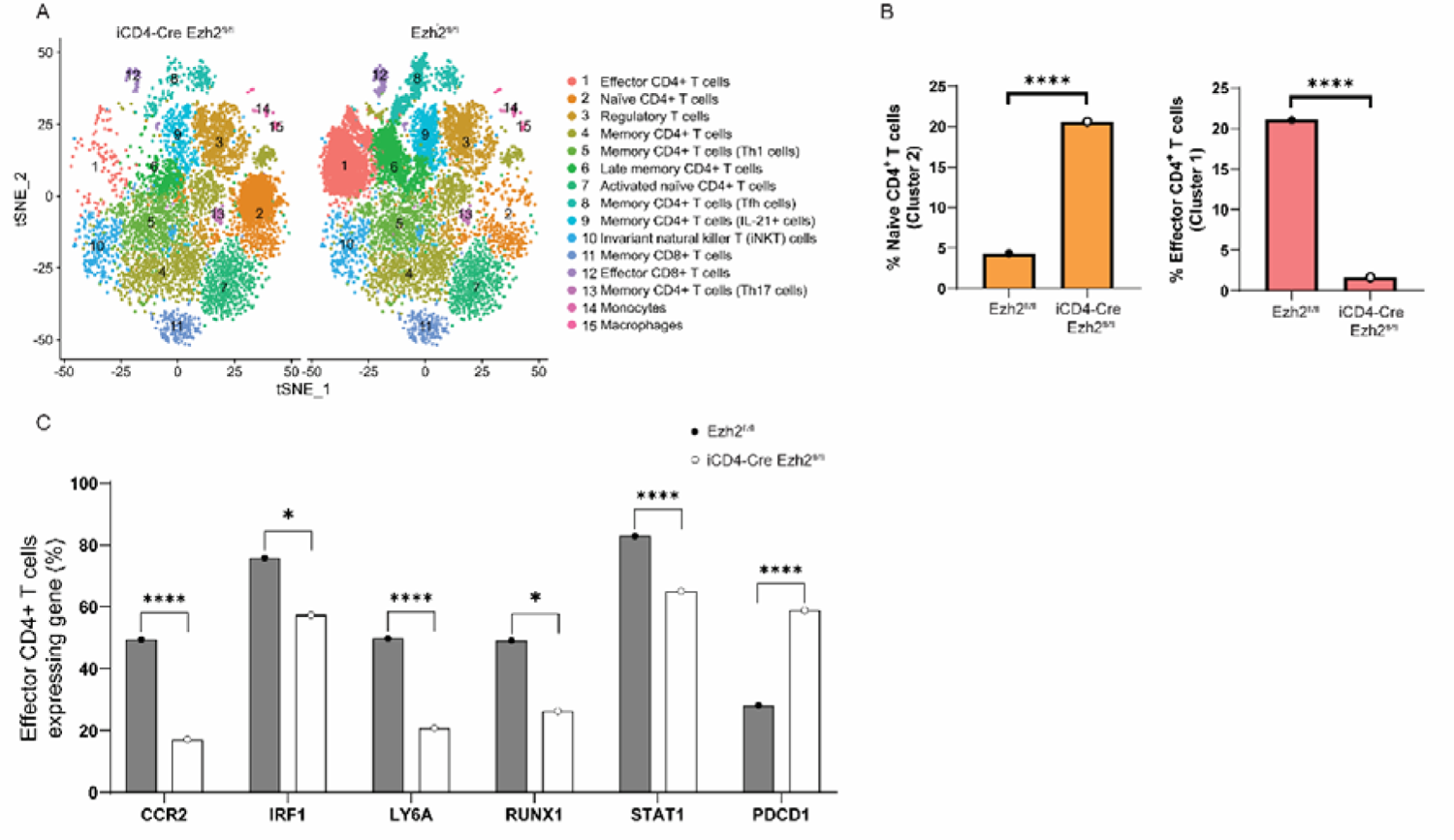
Single-cell RNA-sequencing of CD4+ T cells isolated from the spleen in female *Ezh2^fl/fl^* and iCD4-Cre *Ezh2^fl/fl^* mice from the preventative model. (**A**) tSNE plot of cell clusters identified. (**B**) Frequency of naïve CD4+ T cells and effector CD4+ T cells in *Ezh2^fl/fl^* and iCD4-Cre *Ezh2^fl/fl^* mice. ****p<0.0001, proportion test. (**C**) Proportion of effector CD4+ T cells expressing indicated genes in *Ezh2^fl/fl^* and iCD4-Cre *Ezh2^fl/fl^* mice. \**p*<0.05, ****p<0.0001, proportion test.

We analyzed the proportions of each cell cluster to determine differences in T cell subset composition between iCD4-Cre *Ezh2^fl/fl^* and control *Ezh2^fl/fl^* mice. The most significant differences were observed in the proportion of naïve and effector CD4+ T cells. iCD4-Cre *Ezh2^fl/fl^* mice had significantly more naïve and less effector CD4+ T cells in the spleen (**Figure 5B**). Further, iCD4-Cre *Ezh2^fl/fl^* mice had significantly less late memory CD4+ T cells and follicular helper CD4+ T cells compared to *Ezh2^fl/fl^*mice (**Supplementary** Figure 4).

Our data demonstrate significant impairment in the differentiation of naïve CD4+ T cells into effector CD4+ T cells with EZH2 deficiency. To investigate the effect of EZH2 deficiency on the functional status of effector CD4+ T cells as reflected by their transcriptional profile, we compared gene expression profiles in effector CD4+ T cells between iCD4-Cre *Ezh2^fl/fl^* and control *Ezh2^fl/fl^* mice. In total, we detected 38 differentially expressed genes, of which 26 and 12 genes were upregulated and downregulated in EZH2-deficient T cells, respectively (**Supplementary Table 1**).

Gene Ontology analysis focused on Biological Process revealed that 6 out of 12 downregulated genes belong to “positive regulation of leukocyte differentiation” Gene Ontology (GO:1902107, P value= 3.784E-7) (**Supplementary Table 2**). These include key genes involved in T cell function, such as *RUNX1*, *IRF1*, *JUN, CCR2*, *CCL5*, and *FOS,* which were significantly downregulated in effector CD4+ T cells from iCD4-Cre *Ezh2^fl/fl^* compared to control *Ezh2^fl/fl^* mice. Other genes involved in T cell function such as *STAT1* and *LY6A* were also downregulated in EZH2-deficient mice. The negative T cell regulator gene *PDCD1* was significantly upregulated in effector CD4+ T cells in iCD4-Cre *Ezh2^fl/fl^* compared to control *Ezh2^fl/fl^* mice (**Figure 5C, Supplementary Table 1**). These data suggest that EZH2 deficiency is associated with reduced numbers and impaired function of effector CD4+ T cells.

To understand what might be driving EZH2-dependent impaired CD4+ T cell transition from naïve to effector CD4+ T cells, we compared single-cell transcriptional profiles in naïve CD4+ T cells between iCD4-Cre *Ezh2^fl/fl^* and control *Ezh2^fl/fl^* mice. We detected 77 differentially expressed genes, of which 23 and 54 genes were upregulated and downregulated in EZH2-deficient T cells, respectively (**Supplementary Table 3**). Gene Ontology enrichment analysis revealed enrichment in biological processes involved in protein folding and programed cell death (**Supplementary Table 4**). Further analysis of differentially expressed genes indicates increased entry into the S phase of the cell cycle and suppression of protein transport and key immune-related effector molecules in T cells including cytokine genes such as IFNG and TNF (**Supplementary** Figure 5). Functional annotation also suggested activation of apoptosis (z-score= 2.13, P= 3.15E-14) in EZH2-deficient T cells.

To examine if EZH2 deficiency altered T cell clonal repertoire, we used TCR RNA-seq to identify the number of TCR alpha-beta clones in iCD4-Cre *Ezh2^fl/fl^* and control *Ezh2^fl/fl^* mice. These data reveal a reduction in CD4+ T cell clonal diversity in EZH2-deficient mice, with some clones completely disappearing in iCD4-Cre *Ezh2^fl/fl^* compared to control *Ezh2^fl/fl^* mice (**Figure 6**).

**Figure 6.**
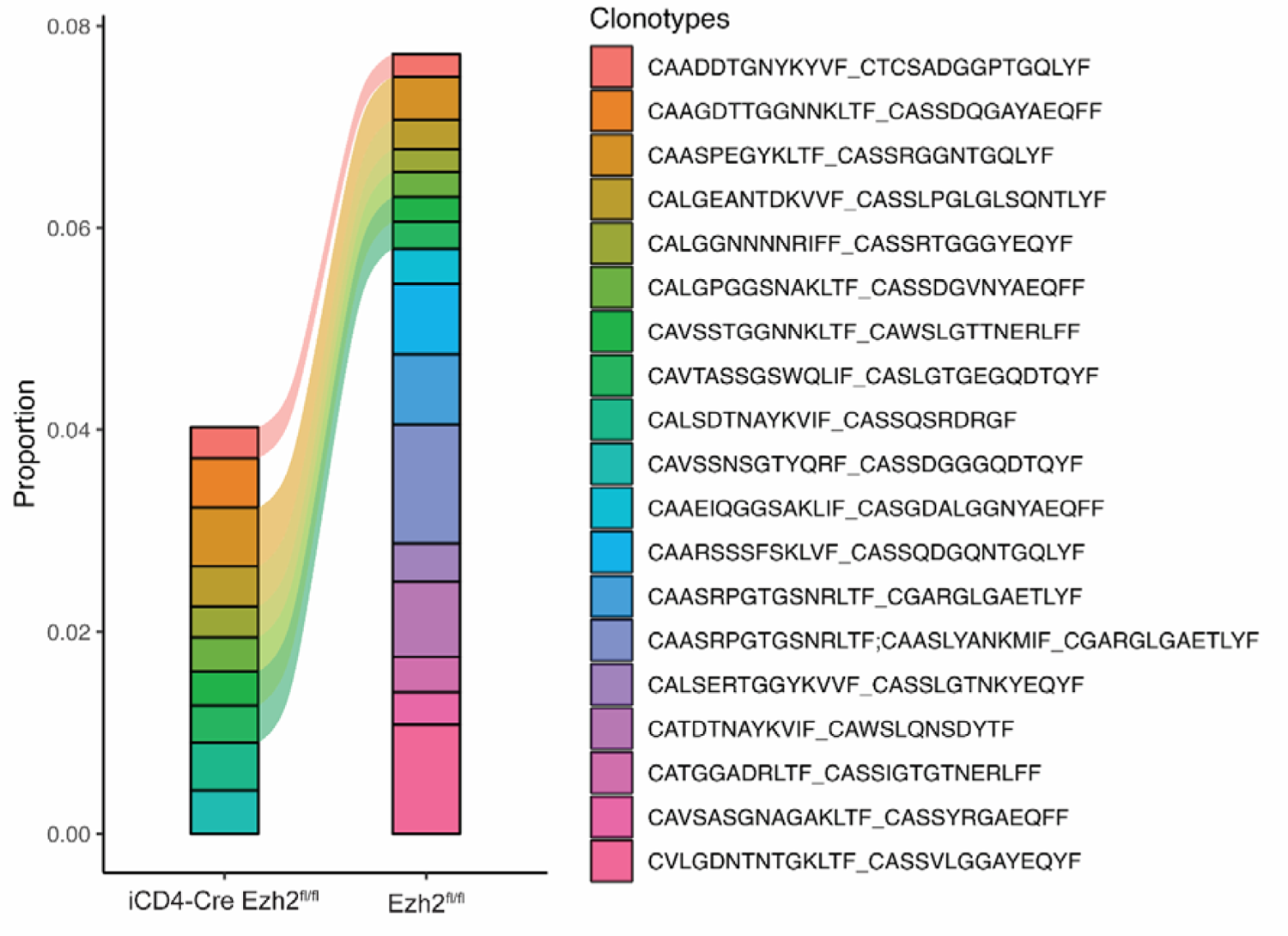
T-cell receptor repertoire analysis in the spleen using single-cell T-cell receptor sequencing in *Ezh2^fl/fl^* and iCD4-Cre *Ezh2^fl/fl^*mice.

### Kidney-infiltrating single-cell RNA-seq reveals T cell subset compositional changes with *Ezh2* deletion

We performed single-cell RNA sequencing in FACS-sorted CD4+ T cells isolated from kidney in iCD4-Cre *Ezh2^fl/fl^* and control *Ezh2^fl/fl^* mice, and identified 12 cell clusters based on transcriptional profiles. Cell identity of each cluster was determined by evaluating differentially expressed genes within the clusters.

These data reveal significant reduction in kidney-infiltrating effector CD4+ T cells in EZH2-deficient mice, consistent with flow cytometry data (**Supplementary** Figure 6). Differential gene expression revealed 27 and 18 genes upregulated and downregulated in EZH2-deficient mice, respectively (**Supplementary Table 5**). Gene Ontology analysis of downregulated genes revealed a number of Biological Process ontologies related to T cell activation and function (**Supplementary Table 6**).

Notably, key genes related to T cell function such as *STAT1*, *CD2*, *CD4*, *CCR2* and *CCR5*, were significantly downregulated in EZH2-deficient kidney-infiltrating effector CD4+ T cells. *PDCD1* was significantly upregulated in kidney-infiltrating effector CD4+ T cells with EZH2 deficiency. Similar to the spleen, CD4+ T cells showed less clonal diversity in EZH2-deficient mice (**Supplementary** Figure 6).

## Discussion

We generated a mouse model that allows for inducible deletion of *Ezh2* in CD4+ T cells on the MRL/*lpr* lupus-prone genetic background. We demonstrated that inducing EZH2-deficient CD4+ T cells is sufficient to abrogate lupus-like disease. Consistent data were observed when EZH2 deficiency was induced before or after the onset of autoimmunity, in a preventative and a therapeutic *in vivo* model, respectively. We observed significant reduction in proteinuria, and autoantibody production in EZH2-deficient compared to littermate control mice. Interestingly, histological examination showed marked improvement in interstitial nephritis in EZH2-deficient mice, but no difference in glomerulonephritis scores between iCD4-Cre *Ezh2^fl/fl^* and control *Ezh2^fl/fl^* mice. This is in contrast to the effects we previously observed when we deleted *Ezh2* in B cells in MRL/*lpr* mice. Indeed, mice with EZH2-deficiency in B cells demonstrated impaired germinal center B cell and plasmablast differentiation, and had significant reduction in proteinuria but with histological improvement in glomerulonephritis and no effect on interstitial nephritis (9). Taken together, these data suggest that EZH2 deficiency induces distinct cell-type specific but complementary effects on renal involvement in MRL/*lpr* mice. While immune-complex deposition and the resulting glomerulonephritis is the whole mark of lupus nephritis, strong evidence suggests that interstitial nephritis is a predictor of progression to end-stage renal disease in lupus patients (10, 11). We observed a significant reduction in kidney-infiltrating effector CD4+ T cells in EZH2-deficient mice. Further, single-cell RNA-seq of kidney-infiltrating CD4+ T cells revealed a transcriptional profile consistent with impaired T cell function in iCD4-Cre *Ezh2^fl/fl^* compared to *Ezh2^fl/fl^* control mice.

Our data revealed that deleting *Ezh2* in CD4+ T cells impairs T cell differentiation and activation in MRL/*lpr* mice. Flow cytometry and single-cell RNA-seq data confirmed significant reduction in effector CD4+ T cells and the accumulation of naïve CD4+ T cells in the spleen of EZH2-deficient compared to control mice. Furthermore, analysis of kidney-infiltrating T cells revealed a significant reduction in effector CD4+ T cells in EZH2-deficient mice. Single-cell RNA-seq of the T-cell receptor suggests a polyclonal effect of EZH2 deficiency, with evidence of reduced numbers of T cell clonotypes and deletion of T cell clones in both the spleen and in kidney-infiltrating CD4+ T cells in iCD4-Cre *Ezh2^fl/fl^* compared to *Ezh2^fl/fl^* control mice. The specificities of the deleted T cell clones and whether they are autoreactive remains to be determined.

EZH2 expression has been shown to be play an important role in T cell development and function (12). Studies in tumor-infiltrating T cells suggests that EZH2-expressing CD4+ T cells are pro-inflammatory and characterized by polyfunctional cytokine expression (13). They are capable of expressing multiple cytokines, including IL-2, interferon-γ, and TNF (13). It has been suggested that EZH2 induces T cell activation by repressing Notch signaling repressor genes *Numb* and *Fbxw7* (13). We did not observe differential expression of *Numb* or *Fbxw7* in iCD4-Cre *Ezh2^fl/fl^* compared to *Ezh2^fl/fl^* control MRL/*lpr* mice. However, single-cell RNA-seq analysis in effector CD4+ T cells in the spleen revealed downregulation of several key genes involved in T cell function in EZH2-deficient mice. These include *RUNX1*, *IRF1*, *JUN, CCR2*, *CCL5*, and *FOS,* which are involved in T cell proliferation, activation, migration, differentiation, and cytokine production (14–18). These data suggest that impaired differentiation and function of effector CD4+ T cells underlies improvement of lupus-like disease in EZH2-deficient mice. Indeed, downregulation of *CCR2*/*CCL5*, which play a role in T cell migration, might explain significantly reduced abundance of effector CD4+ T cells in the kidneys of EZH2-deficient mice.

To gain further insights into possible mechanisms underlying impaired T cell differentiation from naïve to effector CD4+ T cells in EZH2-deficient mice, we compared gene expression profiles in naïve CD4+ T cells between iCD4-Cre *Ezh2^fl/fl^* and control *Ezh2^fl/fl^* mice. Functional bioinformatics analysis suggests that EZH2 deficiency favors the entry of naïve CD4+ T cells into the S phase of the cell cycle and the activation of programed cell death pathways. This is consistent with previous reports suggesting that EZH2 expression in T cells confers resistance to apoptosis (13, 19). Further, the negative T cell regulator gene *PDCD1* was upregulated in effector CD4+ T cells in both the spleen and the kidney in EZH2-deficient mice In summary, our data provide additional support for the role of EZH2 in autoimmunity and the potential therapeutic implications of targeting EZH2 in lupus. Enhancing CD4+ T cell entry into the S phase of the cell cycle, activation of programed cell death, and reduced effector T cell function by suppressing key T cell transcriptional regulators appear to mediate the potential therapeutic effect of inhibiting EZH2 in the setting of autoimmunity. Importantly, T-cell EZH2 deficiency prevents the accumulation of effector CD4+ T cells in the kidneys and the development of tubulointerstitial nephritis which is a strong predictor of end-stage renal disease in patients with lupus nephritis. Further studies to explore the repurposing of EZH2 inhibitors in the setting of clinical trials in lupus nephritis are warranted.

## Methods

### Mice

Female B6(129X1)-Tg(Cd4-cre/ERT2)11Gnri/J mice from the Jackson Laboratory were backcrossed to MRL*/lpr* male mice. A speed congenic method was used by selecting the top two out of 15-20 male pups with the highest MRL*/lpr* genetic background from each generation for subsequent backcrossing. After 6 generations, a mouse reached 99.8% MRL*/lpr* background and was crossed with female *EZH2^fl/fl^* MRL*/lpr* mice we recently generated (9), to get *Ezh2^fl/fl^* and CD4-Cre/ERT2 *Ezh2^fl/fl^* (abbreviated as iCD4-Cre *Ezh2^fl/fl^*) mice. Both groups of mice received oral gavage administration of tamoxifen (300 mg/Kg) dissolved in corn oil 3 days a week (on Mondays, Wednesdays and Fridays), every other week starting from 10 weeks of age in the preventative model and 14 weeks of age in the therapeutic model until mice reached 24 weeks of age. The *Ezh2^fl/fl^* and littermate iCD4-Cre *Ezh2^fl/fl^* mice were housed together in a pathogen-free environment. Upon euthanasia, we isolated superficial and deep cervical lymph nodes, spleens, and kidneys. All animal work has been approved by the University of Pittsburgh Institutional Animal Care and Use Committee (IACUC).

### Plasma autoantibody measurements

Following mouse euthanasia, cardio puncture was performed to collect blood in K2EDTA tubes. Plasma was isolated by centrifuging at 1500 rcf for 15 minutes, and stored at −80°C. We measured plasma levels of anti-doubled-stranded (anti-dsDNA) IgG and anti-Sm antibodies using an ELISA method, following the manufacturer’s instructions (Alpha Diagnostic International, San Antonio, USA).

### Proteinuria and nephritis assessment

Mouse urine was collected less than a week before euthanasia and stored at −80°C. Urine albumin (Alpha Diagnostic International, San Antonio, USA) and creatinine (R&D Systems, Minneapolis, USA) were measured by following the kit instructions. Urine albumin to creatinine ratio (UACR) was used to evaluate proteinuria.

One kidney from each mouse, after capsule removal, was fixed in formalin, and stored in 70% ethanol before paraffin embedding, sectioning, and staining with hematoxylin and eosin (H&E). Kidney H&E stained slides were then reviewed by a renal pathologist in a blinded manner. Inflammation in glomeruli and interstitial areas was quantitively scored as previously described (8).

### Single-cell RNA sequencing (scRNA-seq) and single-cell T-cell receptor sequencing (scTCR-seq) of splenic CD4+ T cells and kidney-infiltrating CD4+ T cells

Spleens were mashed through 70-µm cell strainers with a syringe plunger and rinsed with PBS to prepare into single-cell suspensions. For kidney dissociation, we used the Multi Tissue Dissociation Kit 2 from Miltenyi Biotec and followed the protocol for mouse kidney dissociation. To preserve cell surface epitopes, we decreased the volume of Enzyme P in the dissociation buffer and the kit’s components for each kidney was as follows: 4.8 mL of Buffer X, 5 µL Enzyme P, 50 µL Buffer Y, 100 µL Enzyme D, and 20 µL Enzyme A. Kidney capsules were removed, and kidneys were placed into gentleMACS C tubes (Miltenyi Biotec) with dissociation buffer and quartered. GentleMACS Octo Dissociator with heating function (Miltenyi Biotec) was used and the program ‘37C_Multi_E’ was run to dissociate kidneys. Cell suspension went through a 70-µm cell strainer and was rinsed with PBS. Red blood cells (RBCs) in spleen and kidneys were lysed with RBC lysis buffer (Invitrogen). Cells were resuspended in PBS and stained with LIVE/DEAD fixable near-IR dead cells stain kit (Invitrogen). Then cells were blocked with anti-mouse CD16/CD32 (BD Biosciences) followed by staining with the following antibodies: anti-CD3 epsilon-VioBlue (Clone 17A2, Miltenyi Biotec), anti-TCR β-Alexa Fluor 488 (Clone H57-597, BioLegend), anti-CD4-V500 (Clone RM4-5, BD Biosciences), and anti-CD8a-PE-Cyanine5 (Clone 53-6.7, BioLegend) in PBS with 1% fetal bovine serum (FBS). CD4+ T cells from a total of 10 *Ezh2^fl/fl^* mice and 9 iCD4-Cre *Ezh2^fl/fl^* mice were sorted by FACSAria cell sorter in the Unified Flow Core at the Department of Immunology in the University of Pittsburgh.

CD4+ T cells isolated from a single organ in each mouse were individually incubated with unique TotalSeq-C anti-mouse antibodies from BioLegend. All CD4+ T cells were barcoded in the above manner to distinguish between different organs and mice before pooling cells together. Sequencing libraries of cell surface protein feature barcoding, single cell 5’ gene expression, and paired single-cell T-cell receptor (TCR) were prepared according to instructions provided by10X Genomics in the Single Cell Core at the University of Pittsburgh. These libraries were sequenced at the UPMC Genome Center. In total, 10,271 splenic CD4+ T cells from *Ezh2^fl/fl^* mice were sequenced, with a mean of 26,854 reads per cell, and 7,704 splenic CD4+ T cells from iCD4-Cre *Ezh2^fl/fl^* mice were sequenced with a mean of 22,603 reads per cell.

In the kidneys, 8,850 kidney-infiltrating CD4+ T cells from *Ezh2^fl/fl^* mice were sequenced, with a mean of 24,796 reads per cell, and 4,066 kidney-infiltrating CD4+ T cells from iCD4-Cre *Ezh2^fl/fl^* mice were sequenced with a mean of 25,649 reads per cell.

### Single-cell sequencing data analysis

Cell Ranger v. 7.1.0 (10x Genomics, Pleasanton, CA, USA) was used to process raw sequencing data following the 10x instructions (https://www.10xgenomics.com/analysis-guides/demultiplexing-and-analyzing-5'-immune-profiling-libraries-pooled-with-hashtags). Briefly, gene expression and T-cell receptor data were demultiplexed with ‘cellranger multi’ and organ-specific data were aggregated using ‘cellranger aggr’. Gene expression data analysis was performed using Seurat v.5.0.1 R package following the standard workflow for visualization and clustering (“Seurat - Guided Clustering Tutorial” vignette). Clustering resolution (“resolution” parameter in the FindClusters function) was set to 0.5 to obtain biologically expected number of clusters. Cluster-specific conserved markers were detected using the FindConservedMarkers function.

Clusters were annotated using scTyper v.0.1.0 (20), singleR v.2.4.1 (21), clustermole v.1.1.1 and manually annotated based on cluster-specific differentially expressed genes. Selected clusters were merged to represent major cell types (for kidney data, two clusters were merged into “Activated naïve CD4+ T cells” and three clusters were merged into “Effector CD4+ T cells”; for spleen, two clusters were merged into “Memory CD4+ T cells”). Immune receptor sequencing data for each condition were annotated with cluster-specific barcodes defined at the RNA-seq analysis step and analyzed using the scRepertoire v.1.12.0 R package. Plots were made using Seurat’s visualization functions or the ggplot2 v3.4.4 R package. All computations were performed in R/Bioconductor v.4.3.2.

### Flow cytometry analysis of CD4+ T cells

Single-cell suspension obtained from the spleens and kidneys after RBC lysis was used to characterize CD4+ T cells. LIVE/DEAD fixable near-IR dead cells stain kit (Invitrogen) was used and anti-mouse CD16/CD32 (BD Biosciences) was added to decrease non-specific bindings. The following antibodies were incubated with the cells at 4°C for 30 minutes: anti-CD3 epsilon-VioBlue (Clone 17A2, Miltenyi Biotec), anti-TCR β-Alexa Fluor 488 (Clone H57-597, BioLegend), anti-CD4-V500 (Clone RM4-5, BD Biosciences), anti-CD8a-PE-Cyanine5 (Clone 53-6.7, BioLegend), anti-CD44-Alexa Fluor 647 (Clone IM7, BioLegend), anti-CD62L-PE/Cyanine7 (Clone MEL-14, BioLegend), and anti-279 (PD1)-PE-Dazzle 594 (Clone RMP1-30, BioLegend). Cells were incubated with anti-CD185 (CXCR5)-PerCP/Cyanine5.5 (Clone L138D7, BioLegend) at room temperature for 1 hr, and True-Nuclear Transcription Factor Buffer Set (BioLegend) was used to fix and permeabilize cells before staining with anti-FoxP3-PE (Clone REA788, Miltenyi Biotec). Cell events were recorded with LSRFortessa (BD Biosciences) at the Department of Pediatrics, University of Pittsburgh, and Flowjo (BD Biosciences) was used for data analysis.

### Statistical analysis

Data were presented as either mean ± standard deviation (SD) or median ± interquartile range. Mann-Whitney *U* test was primarily used to compare differences between groups in GraphPad Prism 9.2.0 (GraphPad, San Diego, CA, USA). A *p* value less than 0.05 was considered statistically significant.

## Acknowledgements

This work was supported by the National Institute of Allergy and Infectious Diseases (NIAID) of the National Institutes of Health (NIH) grant number R01 AI097134. We are thankful to Dr. Desiré Casares-Marfil and Dr. Huizhong Long for their help.

## Author contributions statement

XZ: Conducting experiments, acquiring data, analyzing data, contributed to writing the manuscript; MGD: Analyzing data, contributed to writing the manuscript; LE: Conducting experiments, acquiring data. MMB: Conducting experiments, acquiring data; SB: Analyzing data; AHS: Designing research studies, analyzing data, writing the manuscript.

## Notes

### Competing Interest Statement

The authors have declared no competing interest.

### Summary of Updates

Some changes in the results and discussion section to correct errors.

