## Supplementary Figures for "Inducible deletion of *Ezh2* in CD4+ T cells inhibits kidney T cell infiltration and prevents interstitial nephritis in MRL/*lpr* lupus-prone mice"

**Supplementary Figure 1:** Weight of cervical lymph nodes (LN) in *Ezh2^fl/fl^* and iCD4-Cre *Ezh2^fl/fl^* female and male mice from the preventative (**A**) and the therapeutic (**B**) mouse models. Spleen weight in *Ezh2^fl/fl^* and iCD4-Cre *Ezh2^fl/fl^* mice from the preventative (**C**) and the therapeutic (**D**) mouse models. Data are presented as mean ± SD, **p*<0.05, ***p*<0.01, ****p*<0.001, two-tailed Mann-Whitney test.

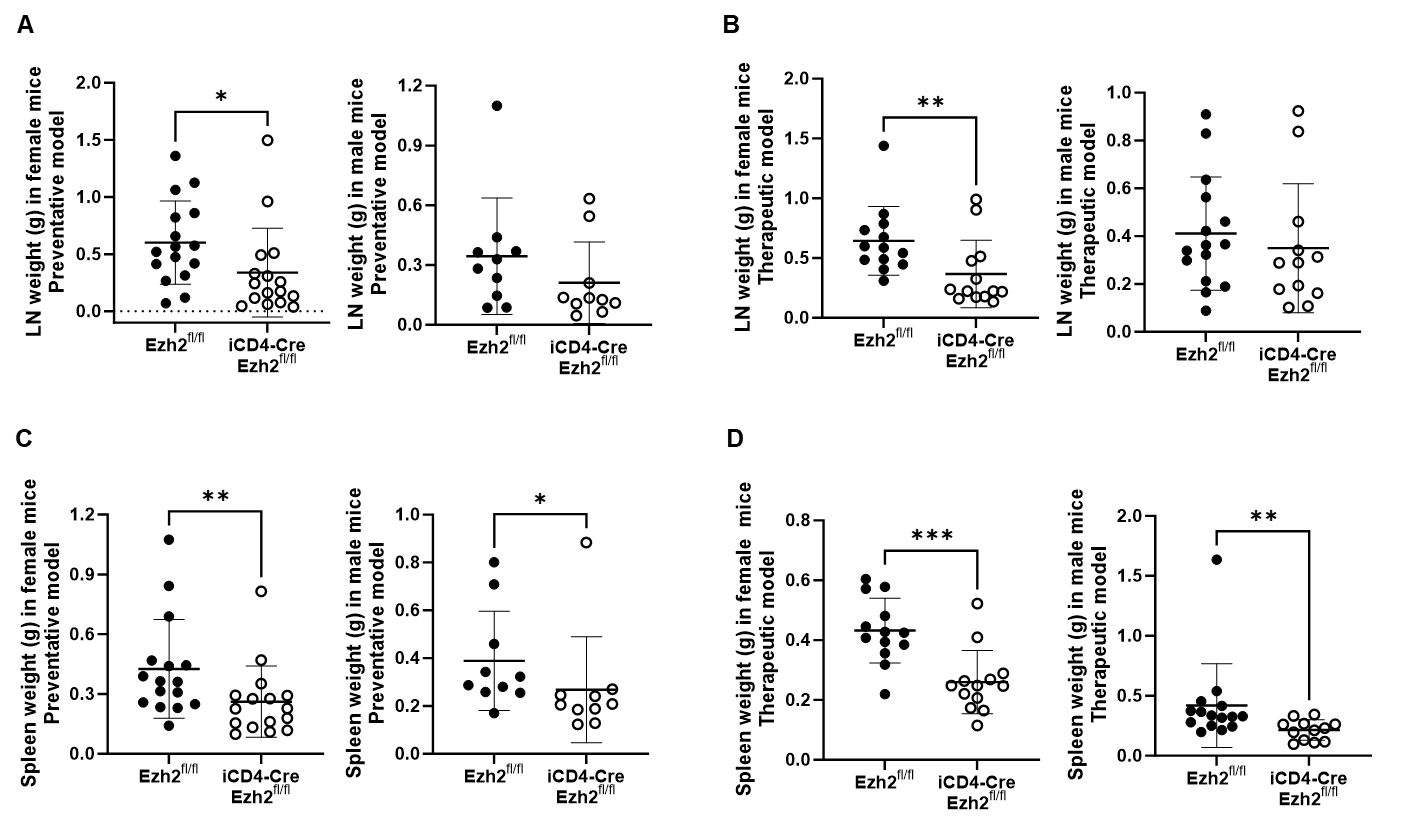

**Supplementary Figure 2:** Flow cytometry gating strategy used to assess T cell subsets in the spleen and kidney of iCD4-Cre *Ezh2^fl/fl^* and *Ezh2^fl/fl^* control mice.

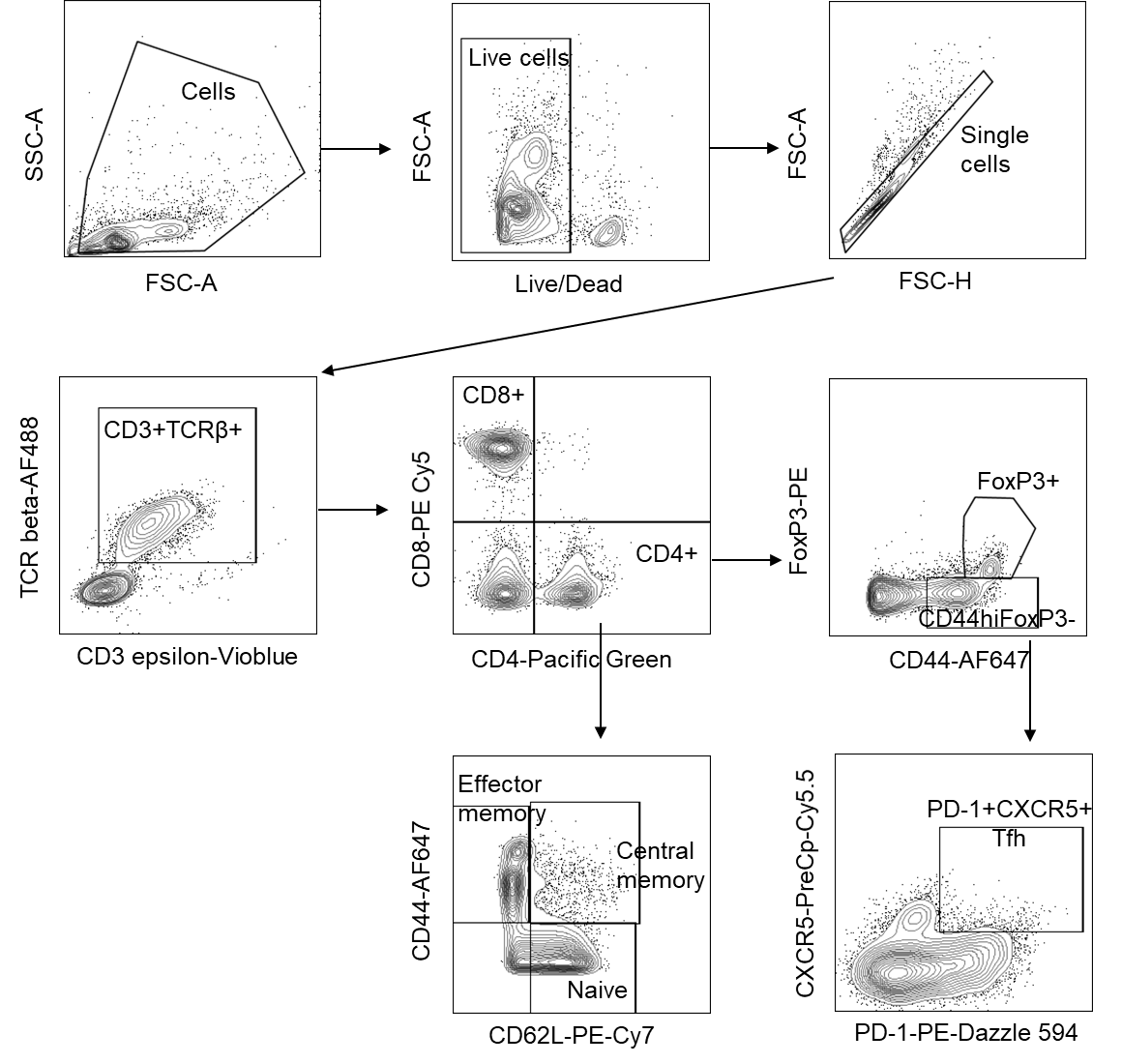

**Supplementary Figure 3:** Statistical analysis of T cell subset numbers and percentages in the spleen of the therapeutic model in *Ezh2^fl/fl^* and iCD4-Cre *Ezh2^fl/fl^* mice. Data are presented as mean ± SD. **p*<0.05, ***p*<0.01, ****p*<0.001, *****p*<0.0001, two-tailed Mann-Whitney test.

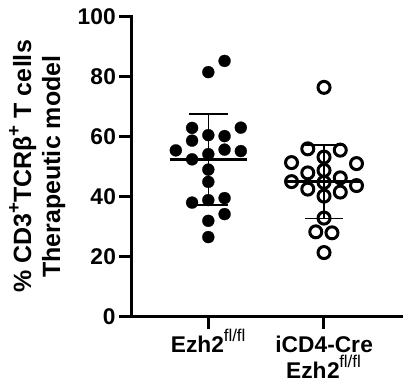

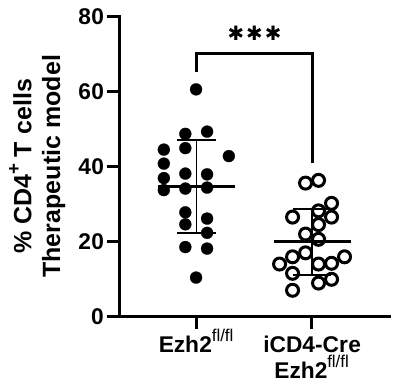

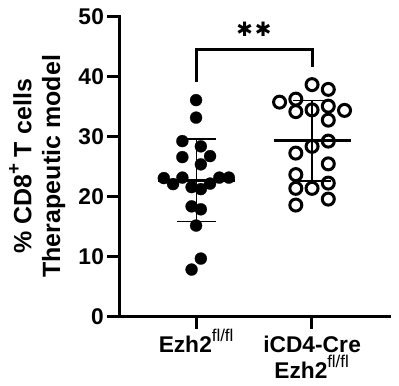

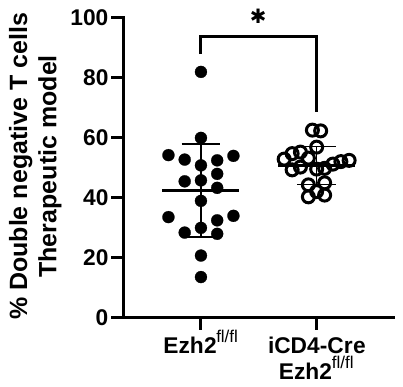

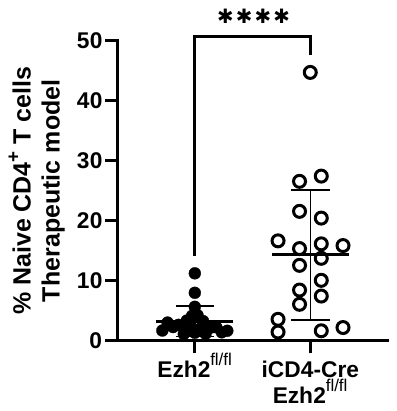

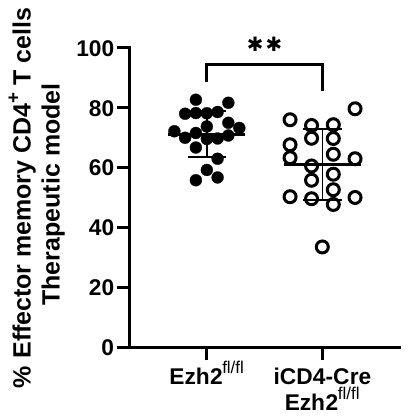

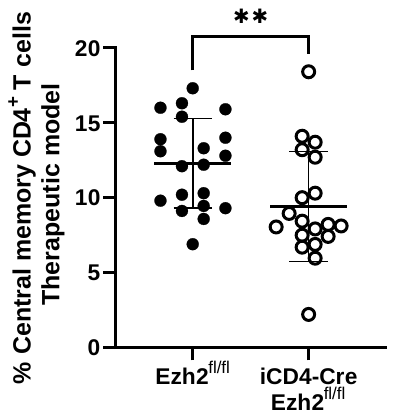

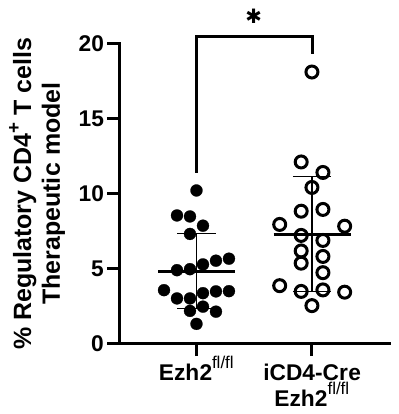

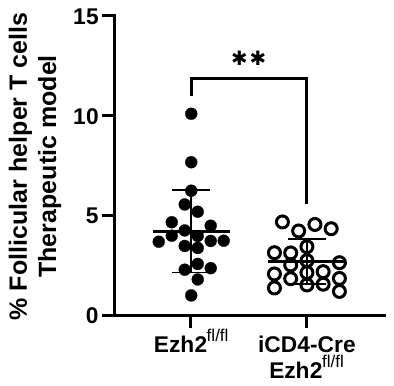

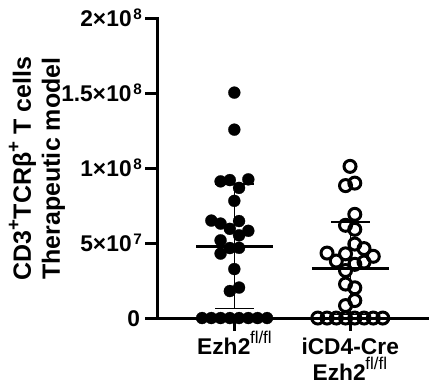

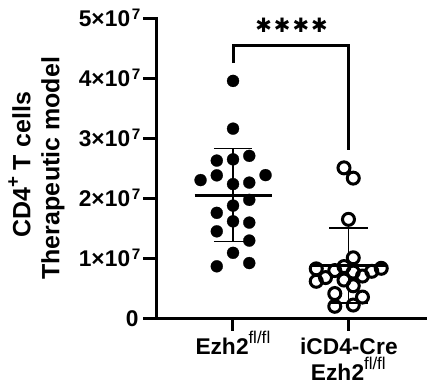

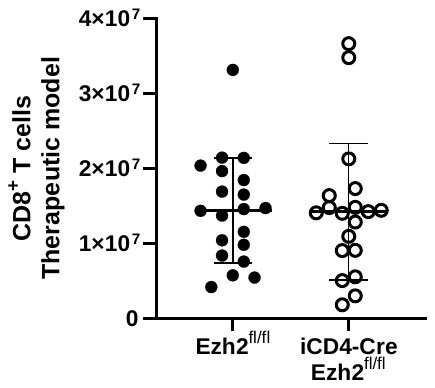

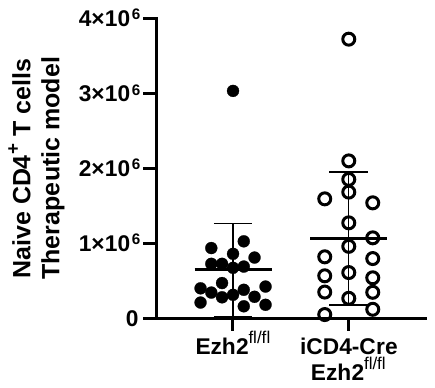

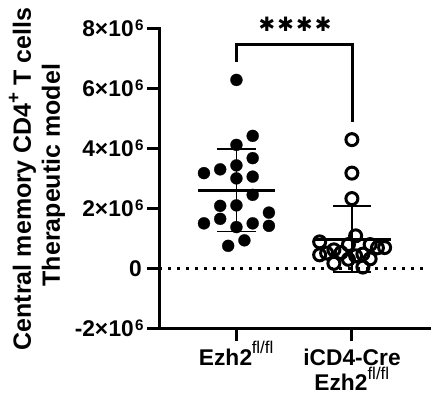

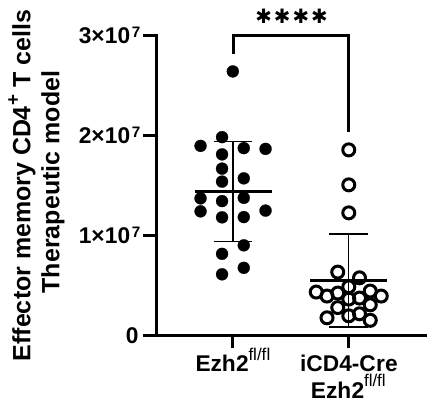

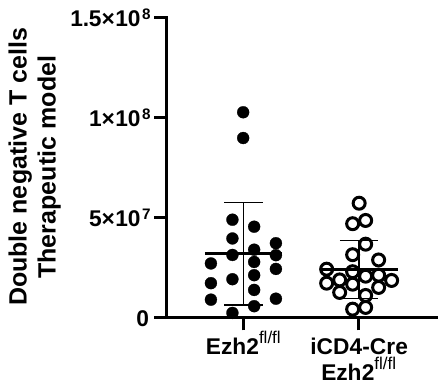

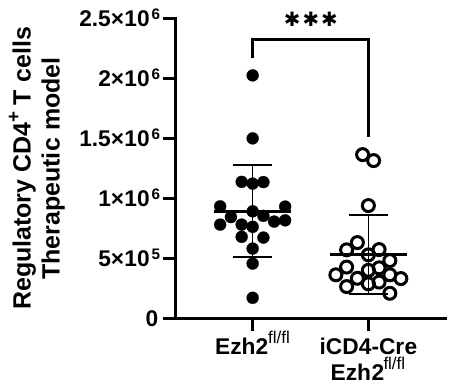

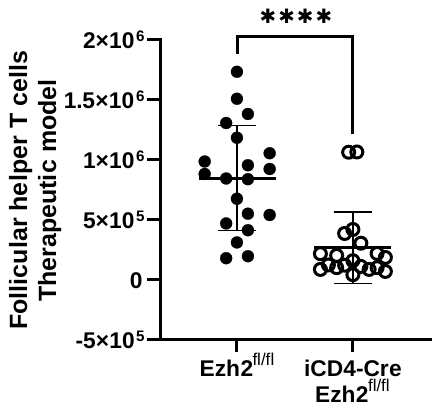

**Supplementary Figure 4:** Single-cell RNA sequencing analysis revealing differences in the proportions of specific T cell clusters in the spleens of *Ezh2^fl/fl^* and iCD4-Cre *Ezh2^fl/fl^* mice. *****p*<0.0001, proportion test.

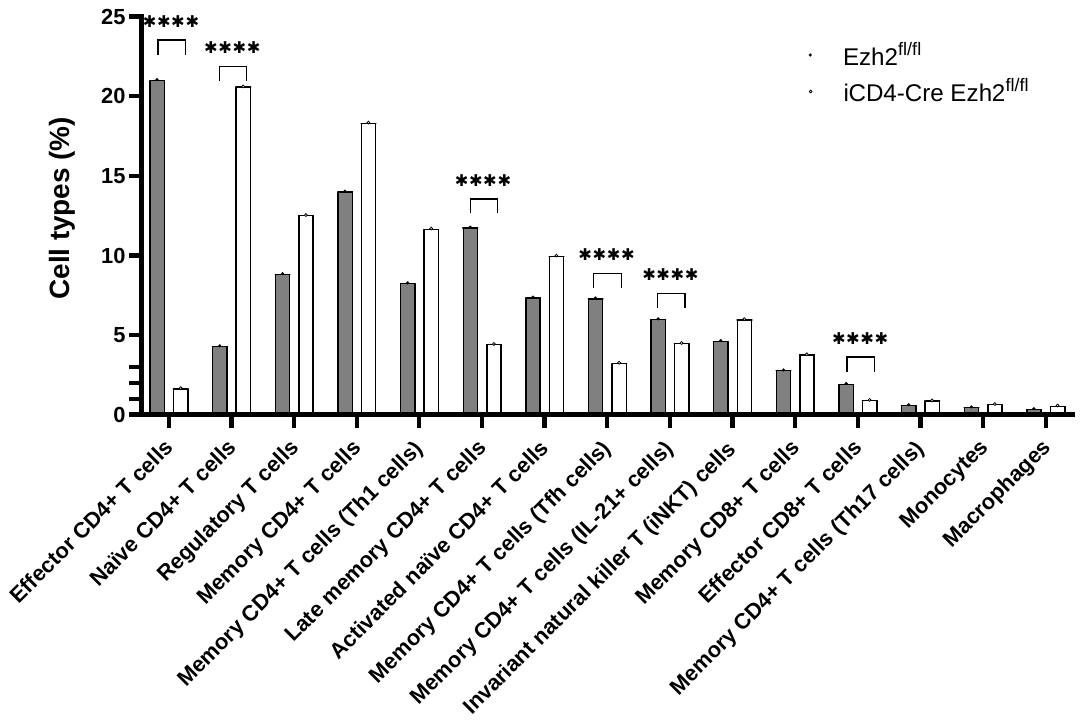

**Supplementary Figure 5:** Graphical summary of the results from the functional enrichment and pathway analysis of genes differentially expressed in naïve CD4+ T cells between iCD4-Cre *Ezh2^fl/fl^* and *Ezh2^fl/fl^* control mice. The analysis was performed using Ingenuity Pathway Analysis (IPA, Qiagen).

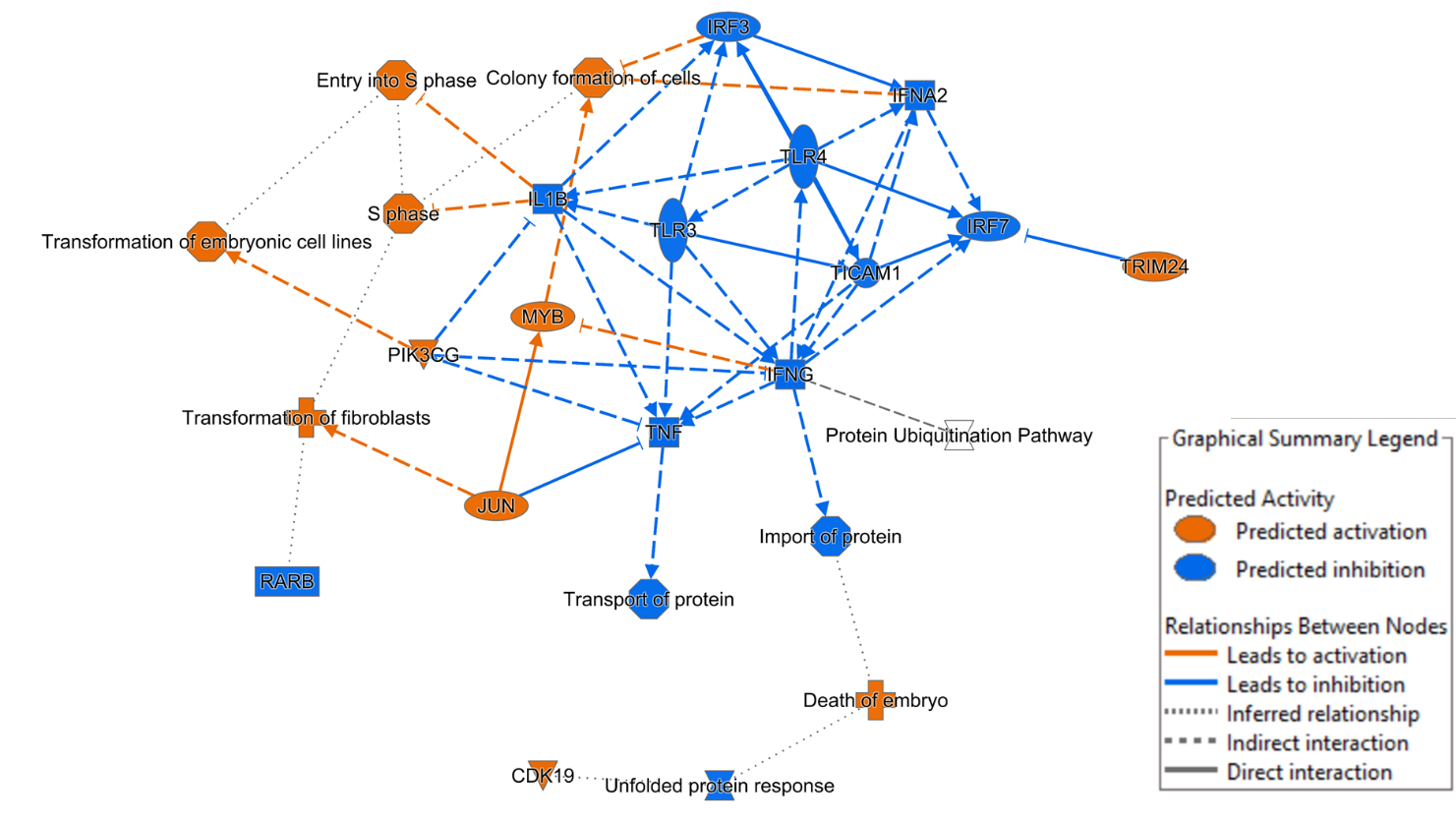

**Supplementary Figure 6:** Single-cell RNA-sequencing of CD4+ T cells isolated from the kidneys of female *Ezh2^fl/fl^* and iCD4-Cre *Ezh2^fl/fl^* mice from the preventative model. (**A**) tSNE plot of cell clusters identified. (**B**) Frequency of effector CD4+ T cells in *Ezh2^fl/fl^* and iCD4-Cre *Ezh2^fl/fl^* mice. ****p<0.0001. (**C**) T-cell receptor repertoire analysis using single-cell T-cell receptor sequencing in *Ezh2^fl/fl^* and iCD4-Cre *Ezh2^fl/fl^* mice.

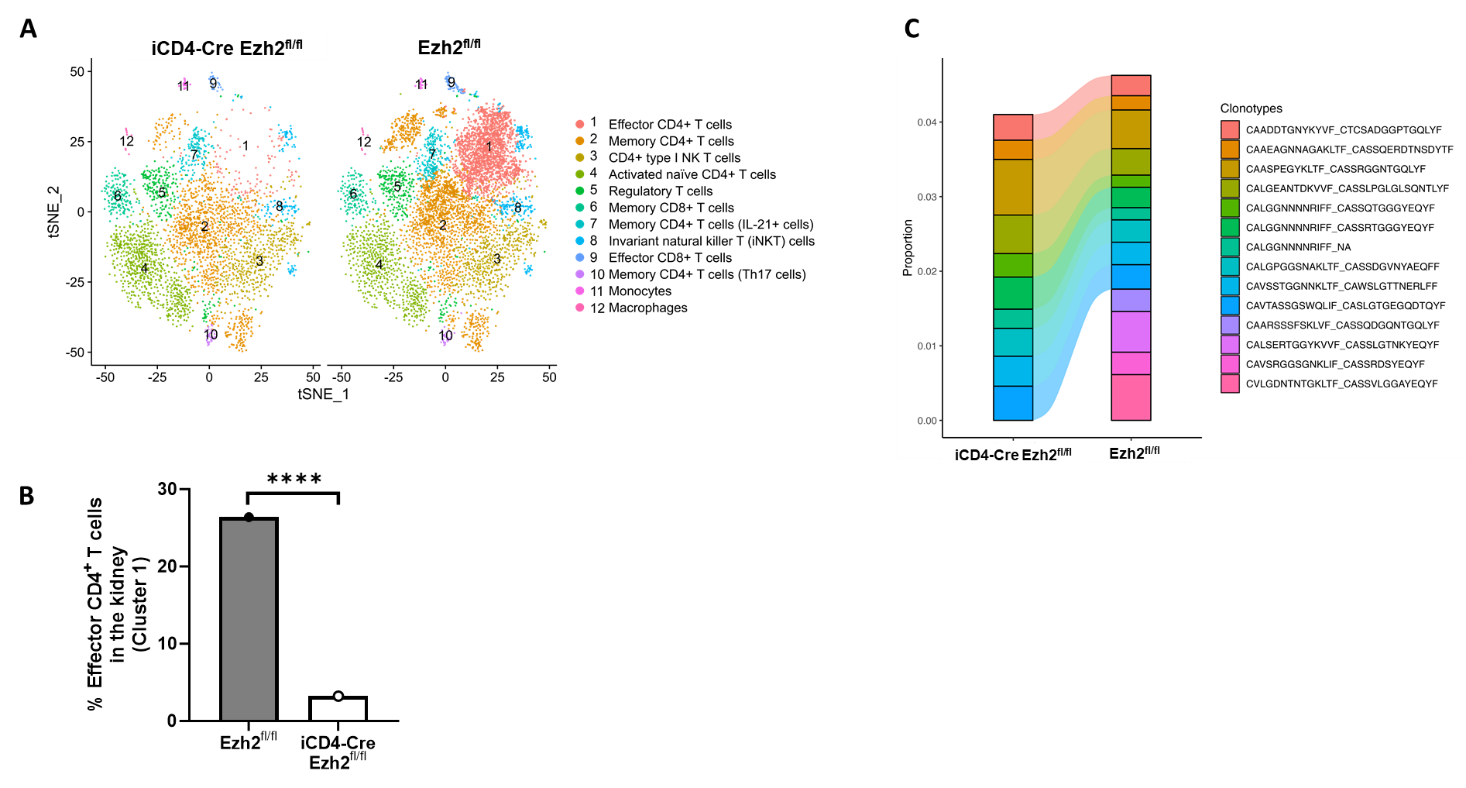
